# Structural Covariance Network Properties Predictive of Early Adolescent Alcohol Initiation

**DOI:** 10.1101/2025.05.08.652983

**Authors:** H. Byrne, R. Visontay, E. K. Devine, N. E. Wade, J. Jacobus, L. M. Squeglia, L. Mewton

## Abstract

**Importance:** Early alcohol initiation (before age 15) is associated with adverse outcomes. Understanding mechanisms behind early alcohol initiation is essential for informing prevention efforts.

**Objective:** To examine whether structural covariance network properties at ages 9-10 years predict early alcohol initiation.

**Design:** Case-control, population-based study design.

**Setting:** Data from the Adolescent Brain Cognitive Development study were used. Baseline structural brain imaging data (ages 9-10) were used for generation and comparison of structural covariance networks. Data from baseline to 4-year follow-up (≤age 15) assessments were used to determine alcohol initiation.

**Participants:** Participants were excluded if they reported consuming a full drink of alcohol at baseline, or did not meet imaging inclusion criteria. Controls were excluded if they had not yet been assessed or were missing substance use data at 4-year follow-up. In total, 3,878 participants met study criteria, of which 182 participants initiated alcohol. Structural covariance network properties were compared between the full sample and a 1:1 propensity-matched sample based on age, sex, race, ethnicity, religion, parental education, prenatal alcohol exposure, and baseline alcohol sipping.

**Main Outcomes and Measures:** Structural covariance networks were estimated using regional cortical thickness and volume measurements. Measures of network segregation (modularity, clustering coefficient), integration (characteristic path length, global efficiency), and resilience (degree assortativity) were compared between groups. Early alcohol initiation was defined as consuming a full drink between baseline and 4-year follow-up

**Results:** Alcohol initiators (*n*=182, median[IQR] age, 10.3[9.9-10.8]; 101 female[55.5%]) demonstrated lower network segregation (modularity: area-under-the-curve[AUC] difference[95%CI]=-0.017[-0.017,-0.007], *p*=0.030; clustering coefficient: AUC[95%CI]=-0.026[-0.027,-0.012], *p*=0.0495) and higher network integration (characteristic path length: AUC[95%CI]=-0.106[-0.099,-0.046], *p*=0.020; global efficiency: AUC[95%CI]=0.011[0.005,0.011], p=0.010), compared to non-initiators (*n*=3,696, median[IQR] age, 9.9[9.4-10.4]; 1750 female[47.4%]) when controlling for age, sex, and mean cortical thickness. Within the matched sample, only differences in network integration were preserved (characteristic path length: AUC[95%CI]=-0.044[-0.032,0.035], *p*=0.010; global efficiency: AUC[95%CI]=0.003[-0.003,0.003], *p*=0.040). There were no differences between full or matched samples when comparing cortical volume structural covariance networks.

**Conclusions and Relevance:** Differences in cortical thickness structural covariance network properties at ages 9-10 predicted alcohol initiation before age 15. These findings suggest cortical thickness network topology may reflect a neuroanatomical risk marker for early alcohol initiation.

**Key points:** *Question:* Do structural covariance network properties at age 9-10 years predict alcohol initiation prior to age 15?

*Findings:* In this case-control study of 3,878 participants, early adolescent alcohol initiators demonstrated differences in cortical thickness network integration and segregation compared to their non-initiating peers.

*Meaning:* Alcohol-naïve adolescents who initiate alcohol use early in life demonstrate differences in structural brain network organization compared to their abstinent peers, which may reflect a neuroanatomical risk marker for early alcohol use.

## Introduction

Globally, approximately 1.34 billion people consume alcohol in harmful amounts, with alcohol use accounting for 1.78 million deaths in 2020^1^. Alcohol use often begins, and noticeably escalates, throughout adolescence^2^. Importantly, early alcohol initiation (typically defined as use before the age of 15^3^) is a significant risk factor for alcohol use disorder later in life^4^, in addition to several other long-term adverse psychosocial and health consequences^5^, including poorer academic performance^6^, increased risk of psychiatric disorders^7^, and cancer^8^. As a result, it is essential to better understand the risk factors and mechanisms that contribute to early alcohol initiation to inform prevention and early intervention efforts.

A growing body of research suggests that differences in brain structure may predict early alcohol initiation in adolescence. For example, a recent study of 9,804 children from the Adolescent Brain Cognitive Development (ABCD) study^9^ identified that morphometric differences at age 9-10 years in multiple regions of the brain, including the prefrontal cortex and parahippocampal gyrus, were associated with alcohol sipping and/or consuming a full drink of alcohol prior to age 15^10^. These findings are supported by previous studies that have identified lower volume in the dorsolateral prefrontal cortex in children who are substance-naïve as predictive of early^11^ and moderate-to-heavy^12,13^ future alcohol use. However, recent work by Green et al.^15^ found that neuroanatomical predictors of early substance use experimentation were distributed diffusely across the brain, rather than localized to prefrontal areas. This inconsistency between studies may be due to a focus on regional markers, despite more contemporary conceptualizations of the brain as a complex system or network^16^. Despite its importance for cognitive, emotional, and behavioral development^17^, the extent to which this brain network-level complexity in alcohol-naïve adolescents predicts early alcohol initiation is currently unknown.

Structural covariance network (SCN) analysis provides a useful tool for probing the structural organization and complexity of brain networks. Based on evidence that brain regions with similar developmental trajectories are closely related in morphometry, this technique captures the covariance in cortical thickness, volume, and/or surface area between brain regions^18^. Studies have shown SCNs are sensitive to neurodevelopmental and age-related changes^19^, and importantly, may be predictive of alcohol use in later life^20^. Specifically, Ottino-González et al.^20^ identified that adolescents who exhibit heavy drinking behaviors at age 19 demonstrate differences in cortical thickness SCN properties at age 14. However, only 37% of these participants were alcohol-naïve at baseline; as recent work has shown SCN properties can be affected by both recent^21^ and heavy^22^ substance use, it is unclear whether differences in structural brain organization exist prior to alcohol use, and importantly, whether such features are predictive of early alcohol initiation.

Using data from the ABCD study, this work aimed to explore whether SCN properties in alcohol-naïve 9–10-year-olds are predictive of early initiation of alcohol use (defined as first full drink prior to age 15). We first examined whether regional cortical thickness and volume were predictive of early alcohol initiation. Next, we investigated whether whole brain SCN properties derived from cortical thickness and volume measurements at ages 9-10 differed between early initiators and abstainers.

## Methods

### Participants

Data from the ABCD Study release 5.1 were used, which includes biological and behavioral data from 11,868 participants aged 9-10 years at baseline^9^. Briefly, participants were recruited from 21 sites across the United States between September 2016 and November 2018, with follow-up visits conducted at yearly intervals. Parents/caregivers provided signed informed consent and all participants gave assent. The ABCD protocol was approved by the centralised institutional review board (IRB) at the University of California, San Diego and by the IRBs at the 21 sites.

Release 5.1 includes complete data up to the 3-year follow-up (N=11,868, ages 12-13) and a subset of data for the 4-year follow-up (N=4,754, ages 13-14), due to ongoing data collection during this period. Given the small number of participants who initiated alcohol use at follow-up 3^23^, the current study included data from the subset of participants at follow-up 4. Participants were excluded if they reported consuming a full drink at baseline (n=21), had missing structural MRI data at baseline (n=140), and/or did not meet neuroimaging quality control inclusion criteria (n=458). Participants were excluded if they did not have complete alcohol use data at follow-up 4 or had not yet participated at this timepoint (n=6,708). Finally, because the ABCD dataset is oversampled for siblings and twins, siblings were excluded at random in the final dataset to reduce potential genetic and/or environmental biases (n=663). Of the 11,868 youths enrolled in the study at baseline, 3,878 met our inclusion criteria (*Figure 1*).

**Fig. 1.**
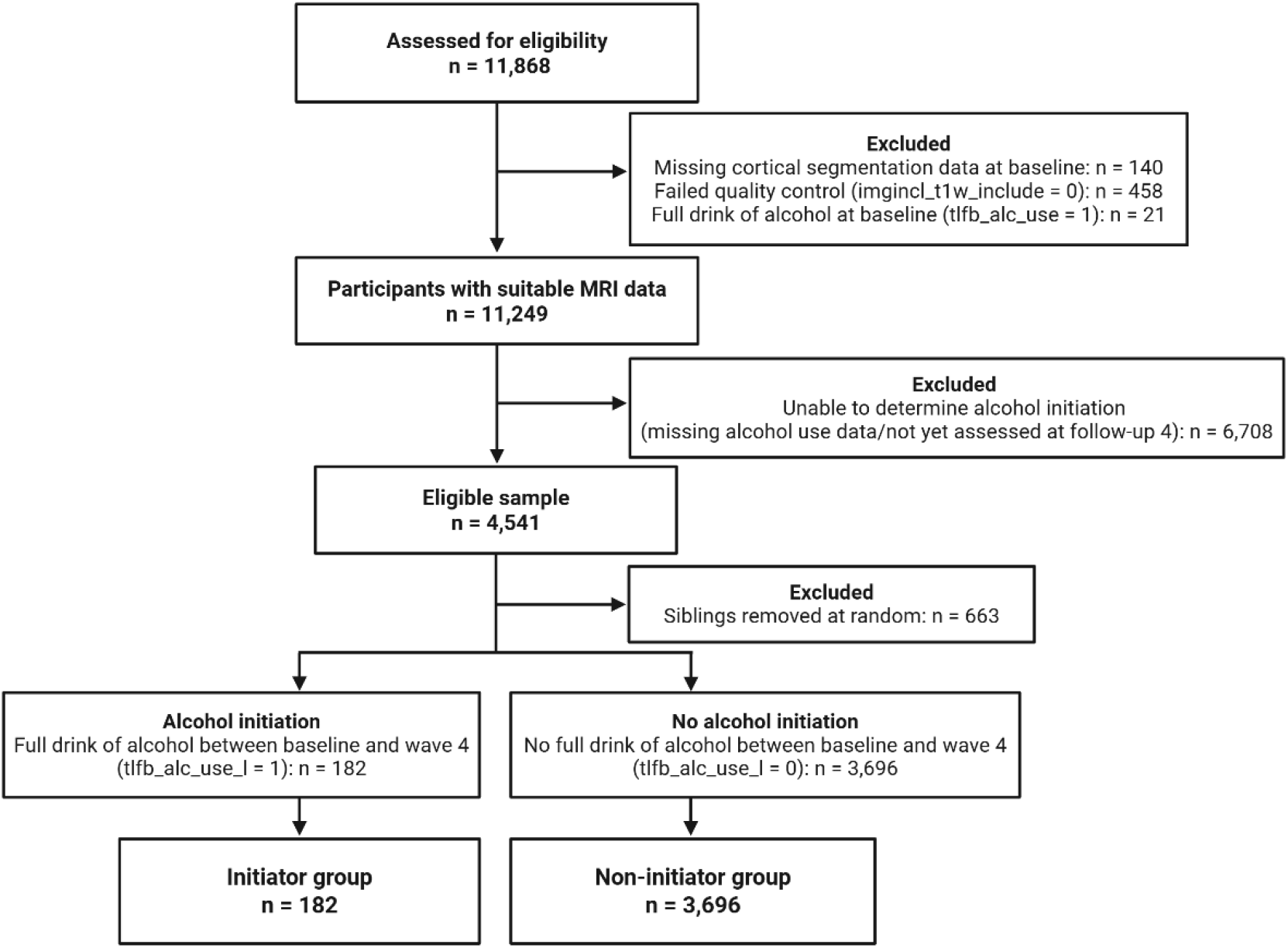
Flowchart for participant inclusion/exclusion criteria. Figure created in *Biorender.com*.

### Measures

#### Alcohol Initiation

Early alcohol use was defined as consuming a full drink of alcohol at any timepoint between baseline and 4-year follow-up (“initiator” group). This was assessed using binary ‘Yes’/’No’ participant responses to the question: “Have you ever tried alcohol at any time in your life? A full drink of beer, wine or liquor (rum, vodka, gin, whiskey),” as self-reported during the Substance Use Interview^24^.

#### Imaging Acquisition

Baseline preprocessed imaging data acquired at ages 9-10 years were used in the current study^25^. Cortical reconstruction and volumetric segmentation of the structural T1-weighted data were performed using *FreeSurfer* (v7.1.1; https://surfer.nmr.mgh.harvard.edu/). Only cortical surface reconstruction data that met recommended quality control criteria were included^25^. Postprocessed cortical thickness and volume measurements mapped to the Desikan-Killiany atlas included 34 cortical parcellations per hemisphere, for a total of 68 regions of interest (ROIs), respectively^26^.

#### SCN generation

All processing and statistical analyses were performed using R Statistical Software (v4.3.3)^27^, conducted in RStudio (v2024.04.2)^28^. First, to minimize scanner effects, imaging data were harmonized prior to analysis using the *RELIEF* package (v0.1.0)^29^. SCNs were generated (*Figure 2*) using the *brainGraph* package (v3.1.0)^30,31^. ROIs for cortical thickness and volume were used to define the ‘nodes’ of the networks. ‘Edges’ were defined as the similarity between pairs of ROIs, calculated using Pearson’s correlations between studentized residuals for each ROI within a given modality. Analyses examining cortical thickness and cortical volume SCNs were conducted separately (generating 68x68 adjacency matrices, respectively).

**Fig. 2.**
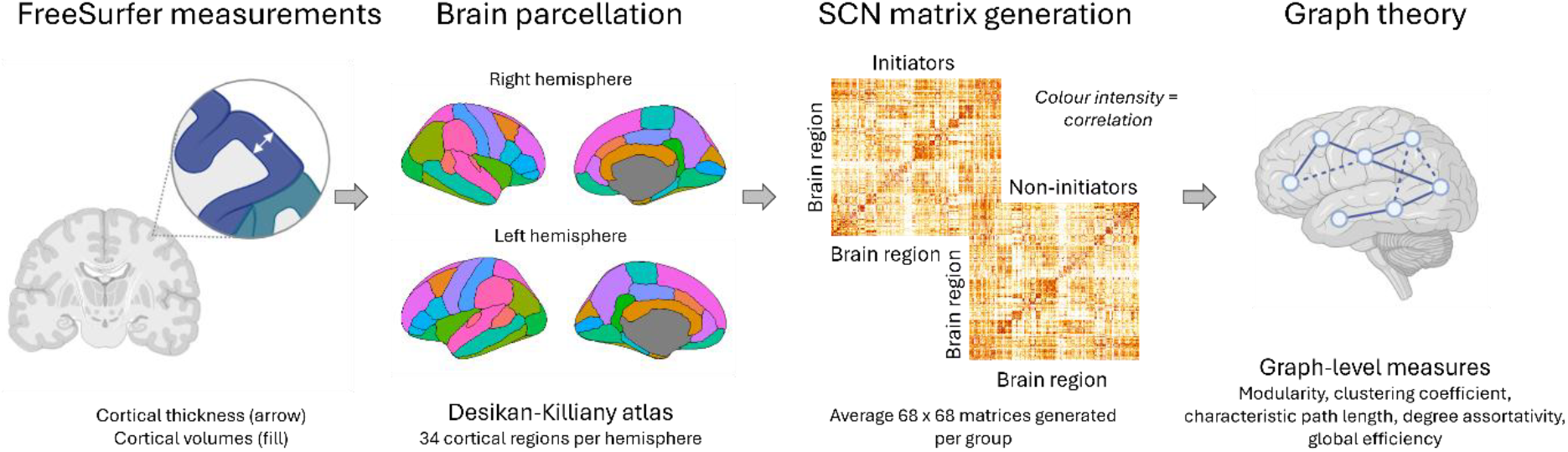
SCN generation pipeline, partially generated using *BioRender.com*.

#### Network measures

Global (whole brain) SCN properties were summarized using graph theory metrics to assess differences in network segregation (modularity and clustering coefficient), integration (characteristic path length and global efficiency), and resilience (degree assortativity) between groups across densities (i.e. the proportion of non-zero edges retained after thresholding). Briefly,^32^ network segregation describes functional specialization of the brain into modules, enabling efficient processing of specific types of information (such as visual input); integration measures the brain’s ability to combine information across regions, enabling coordinated functions such as cognitive processing; resilience refers to the brain’s capacity to maintain function/integrity despite disruptions to the network. These measures have been described in further detail previously^32^ and are summarized in *eTable 1*.

#### Covariates

Current recommendations for neuroimaging analysis in population-based datasets suggest adjusting for age, sex, and global brain measures prior to the inclusion of additional (i.e. sociodemographic) covariates^33–35^. However, unlike the established relationship between regional cortical volume and whole brain volume^36^, there is ongoing debate concerning whether adjusting for mean cortical thickness in regional cortical thickness measurements can obscure meaningful individual heterogeneity^33,37^. Therefore, preliminary testing was conducted to explore the impact of mean cortical thickness on model variance (*Supplementary Materials*; *eTable 3*). This testing confirmed that adjusting for average cortical thickness alongside age and sex in our cortical thickness models was appropriate for our study, with volumetric comparisons adjusted for age, sex, and brain volume (estimated total intracranial volume [eTIV]) as recommended.

Next, to capture the sociodemographic variability in the ABCD sample and to balance sample sizes between groups, the analysis was repeated on a sample matched by first-(age and sex only) and second-level covariates. The second-level covariates chosen for this analysis were identified based on potential relationships with both brain structure and earlier initiation of alcohol use in the ABCD sample. This included race^38^, ethnicity^39^, parental education (highest obtained)^40^, religion^15^, maternal alcohol use during pregnancy^41^ and participant-reported sipping at baseline^42^. Participants were matched at a 1:1 ratio (see *Supplementary Materials*; *eFigure 1*) using the *MatchIt* package (v4.5.5)^43^. Participants were not matched by global brain measures (i.e. mean cortical thickness or eTIV) as the relevance of these measures depend on the measurement being analyzed; therefore, these were adjusted for as a covariate in their respective models in *brainGraph*.

## Statistical Analysis

### Differences in baseline structural brain measurements between groups

To determine whether baseline cortical thickness and volume measurements were associated with initiation of alcohol use within a univariate ROI approach, separate linear regression models were performed with each of the 68 brain regions as outcomes using the *lme4* package (v1.1.35.4)^44^. Models including cortical thickness predictors were adjusted for age, sex, and mean cortical thickness, while models including cortical volume predictors were adjusted for age, sex and eTIV. False Discovery Rate (FDR) correction was used to adjust for multiple comparisons^45^, with statistical significance set to *p*_FDR_<0.05.

### Differences in network measures between groups

To assess differences in structural covariance, permutation testing was used to generate a null distribution of *t*-values for graph measures of network segregation, integration, and resilience between groups for the cortical thickness and volume models. To ensure comparisons were independent of density thresholds, area-under-the-curve (AUC) tests were conducted across 16 densities (0.05–0.2), where each curve represents changes in a specific network measure for each group as a function of network density. To determine the significance of between-group differences in AUC, the observed AUC difference was compared to the corresponding permutation distribution, with *p*-values derived from its percentile position^30^. The groups were shuffled 1000 times, and AUC tests were performed at a two-tailed α of 0.05. AUC results are reported as the primary findings.

### Sensitivity analyses

Two sensitivity analyses were performed. First, to investigate the impact of the second-level covariates in the full sample, a sensitivity analysis was performed repeating the cortical thickness SCN analysis in the full sample using first- and second-level covariates. Second, as individuals who initiated use of other substances during follow-up may present different network properties at baseline compared to total abstainers, a sensitivity analysis was performed excluding participants in the non-initiator group who initiated use of other substances (defined here as cigarette, e-cigarette, or cannabis use) for cortical thickness SCN comparisons (n=142). These analyses are reported in the *Supplementary Materials*.

## Results

### Study sample

Participants who initiated alcohol use (n=182) at any point during follow-up were, on average, older at baseline than those who did not (n=3,696; *Table 1*). A higher proportion of initiators were female, had been exposed to alcohol *in utero*, and reported sipping alcohol at baseline. While differences in race were observed between the groups, no significant differences were found in ethnicity, religion, or parental education. At follow-up, those who initiated alcohol use also reported higher rates of other substances compared to non-initiators. Demographics for the matched sample are reported in *eTable 2*.

**Table 1.** Participant characteristics between early adolescent alcohol initiators and non-initiators.

| Characteristic | Initiators (n = 182) | Non-Initiators (n = 3696) | p-value |
| --- | --- | --- | --- |
| **Baseline** |  |  |  |
| Age at baseline (years)<sup>1,3</sup> | 10.3 (9.9-10.8) | 9.9 (9.4-10.4) | <0.001** |
| Sex<sup>2,4</sup> |  |  | 0.032* |
| Female | 101 (55.5%) | 1750 (47.4%) |  |
| Race<sup>2,5</sup> |  |  | 0.001** |
| Asian | 0 | 105 (2.8%) |  |
| Black/African American, Other/Multiracial<sup>†</sup> | 42 (23%) | 1017 (27.4%) |  |
| White | 140 (76.9%) | 2551 (68.5%) |  |
| Missing/Unknown | 0 | 50 (1.4%) |  |
| Ethnicity<sup>2,4</sup> |  |  | 0.7 |
| Hispanic/Latino | 37 (20.3%) | 798 (21.6%) |  |
| Unknown | 2 (1.1%) | 46 (1.2%) |  |
| Religion<sup>2,5</sup> |  |  | 0.2 |
| Christian | 104 (57.1%) | 2362 (63.9%) |  |
| Jewish, Muslim, Buddhist, Hindu, Something else<sup>†</sup> | 11 (6%) | 188 (5.1%) |  |
| Atheist, Agnostic<sup>†</sup> | 14 (7.7%) | 189 (5.1%) |  |
| Nothing | 47 (25.8%) | 817 (22.1%) |  |
| Missing/Unknown | 6 (3.3%) | 140 (3.78%) |  |
| Highest Parental Education<sup>2,4</sup> |  |  | 0.2 |
| High School | 23 (12.6%) | 549 (14.9%) |  |
| College/Associate's Degree | 63 (34.6%) | 1035 (28%) |  |
| Bachelor's Degree | 43 (23.6%) | 1120 (30.3%) |  |
| Master's Degree | 40 (22%) | 741 (20%) |  |
| Professional/Doctoral Degree | 13 (7.1%) | 246 (6.7%) |  |
| Missing/Unknown | 0 | 5 (13.5%) |  |
| Prenatal alcohol exposure<sup>2,4</sup> |  |  | 0.004** |
| Yes | 62 (34.1%) | 926 (25.1%) |  |
| Missing/Unknown | 13 (7.1%) | 230 (6.2%) |  |
| Alcohol sipping at baseline<sup>2,4</sup> |  |  | <0.001** |
| Yes | 100 (54.9%) | 843 (22.8%) |  |
| Missing/Unknown | 1 (0.5%) | 106 (2.9%) |  |
| **Follow-up** |  |  |  |
| Other substance use (more than a puff)<sup>5</sup> |  |  |  |
| Tobacco cigarette | 24 (13%) | 13 (0.4%) | <0.001** |
| E-cigarette, vape pen, and/or e-hookah | 85 (47%) | 121 (3.3%) | <0.001** |
| Cannabis cigarette (joint) | 56 (31%) | 49 (1.3%) | <0.001** |

<sup>1</sup> Median (IQR); <sup>2</sup> n (%); <sup>3</sup> Wilcoxon rank sum test; <sup>4</sup> Pearson's Chi-squared test; <sup>5</sup> Fisher's exact test. \* = < 0.05; \*\* = < 0.005. <sup>†</sup> Categories have been collapsed for reporting purposes due to low sample sizes and possible identifiability. However, analyses were conducted with all levels of each category preserved.

### Group differences in regional structural brain measurements

After adjusting for age, sex, and mean global cortical thickness, comparisons of cortical thickness between groups revealed differences in the left medial orbitofrontal gyrus and right supramarginal gyrus. However, these differences did not survive FDR correction (*eTable 4*). Similarly, when comparing volumetric measurements, we observed between-group differences in the bilateral superior temporal gyrus and right frontal pole, but these also did not survive FDR correction. No group differences were found in any other ROIs.

### Group differences in cortical thickness structural covariance

When examining group differences in cortical thickness SCN measures adjusting for first-level covariates, the initiator group exhibited lower modularity, clustering coefficient, characteristic path length, and degree assortativity, in addition to higher global efficiency, compared to non-initiators (*Table 2*). When comparing between the matched samples, initiators exhibited lower characteristic path length and degree assortativity, in addition to higher global efficiency, compared to non-initiators. No other differences between groups were observed. These findings were replicated when adjusting for second-level covariates in the full (unmatched) sample, as well as when excluding participants who had used other substances from the alcohol non-initiator group (see *Supplementary materials*). Given the small number of participants initiating alcohol use, further robustness checks were conducted and reported in *Supplementary Materials (eFigures 2 and 3)*.

**Table 2.**
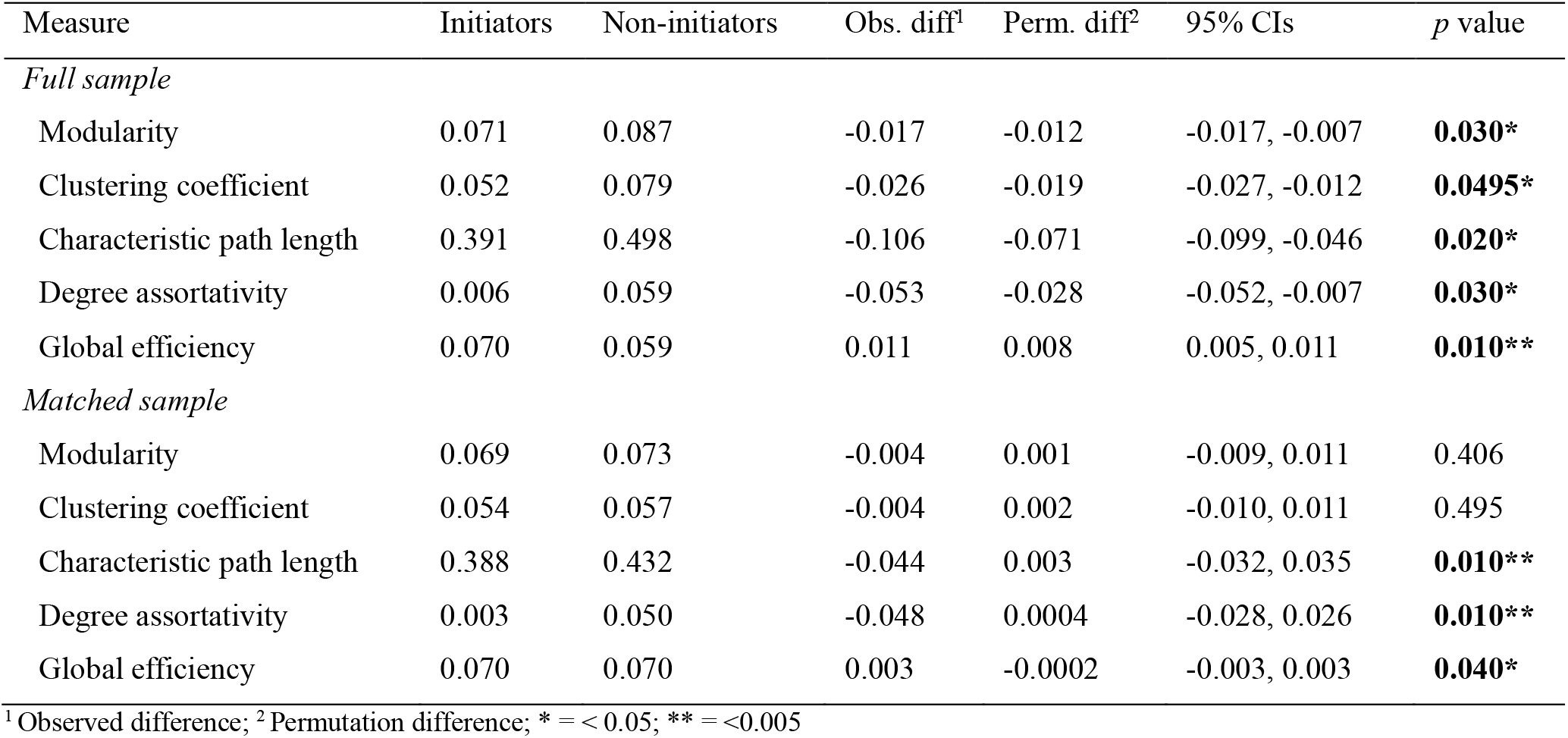
Results from the cortical thickness SCN graph-level AUC network comparisons in the full and matched samples.

### Group differences in cortical volume structural covariance

When controlling for first-level covariates and when comparing against the matched sample, no statistically significant differences in cortical volume SCN measures between groups were identified (*Table 3*). However, similar trends of lower integration and higher segregation in the initiator group were observed. These findings were replicated when adjusting for second-level covariates in the full (unmatched) sample (*Supplementary materials*).

**Table 3.** Results from the cortical volume SCN graph-level AUC network comparisons in the full and matched samples.

| Measure | Initiators | Non-initiators | Obs. diff | Perm. diff | 95% CIs | *p* value |
| --- | --- | --- | --- | --- | --- | --- |
| *Full sample* |  |  |  |  |  |  |
| Modularity | 0.070 | 0.082 | -0.014 | -0.013 | -0.020, -0.007 | 0.436 |
| Clustering coefficient | 0.063 | 0.087 | -0.024 | -0.020 | -0.028, -0.012 | 0.198 |
| Characteristic path length | 0.407 | 0.469 | -0.062 | -0.066 | -0.088, -0.043 | 0.624 |
| Degree assortativity | 0.026 | 0.028 | -0.002 | -0.003 | -0.019, 0.016 | 0.842 |
| Global efficiency | 0.067 | 0.062 | 0.005 | 0.005 | 0.002, 0.007 | 0.446 |
| *Matched sample* |  |  |  |  |  |  |
| Modularity | 0.064 | 0.072 | -0.008 | -0.0004 | -0.010, 0.011 | 0.178 |
| Clustering coefficient | 0.063 | 0.064 | -0.001 | -0.001 | -0.011, 0.009 | 0.911 |
| Characteristic path length | 0.418 | 0.400 | -0.018 | -0.001 | -0.032, 0.025 | 0.267 |
| Degree assortativity | 0.027 | 0.017 | 0.010 | -0.001 | -0.024, 0.019 | 0.485 |
| Global efficiency | 0.067 | 0.067 | 0.0002 | 0.0002 | -0.004, 0.004 | 0.950 |

<sup>1</sup> Observed difference; <sup>2</sup> Permutation difference; \* = < 0.05; \*\* = <0.005

## Discussion

The present study investigated whether whole brain network-level properties in alcohol-naïve children aged 9-10 are predictive of alcohol initiation prior to age 15. Our findings suggest early adolescent alcohol initiators exhibit distinct cortical thickness SCN characteristics, including lower segregation (lower modularity and clustering coefficient) and higher integration (lower characteristic path length and higher global efficiency) compared to non-initiators. These differences were not observed at the regional cortical level, and differences in network integration remained robust after controlling for confounders using a matched sample. However, no similar differences were found in the cortical volume network comparisons, possibly related to the reduced sensitivity of composite measurements^46^. Collectively, these findings provide novel evidence to suggest global brain network organization may play a role in early alcohol use vulnerability, highlighting the need to investigate early life mechanisms driving these differences in brain network development.

The observations made in this study replicate a SCN profile previously identified in adults with alcohol use disorder^20,22^, and in adolescents who later engaged in hazardous drinking^20^. Specifically, Ottino-Gonzalez et al.^20^ identified that lower network segregation and higher integration aged 14 predicted hazardous drinking behaviors at age 19, attributed to delayed maturation of cortico-cortical networks involved in reward- and risk-taking behaviours. However, as network differences between groups were eradicated once excluding alcohol-naïve participants, it is difficult to eliminate the possibility of reverse causation. Our findings are the first to identify that these replicable network differences precede the onset of alcohol use independent of prior alcohol consumption, representing a potential early neuroanatomical marker of future alcohol involvement. While the functional implications of these findings require further investigation, they highlight the potential neurobiological involvement in alcohol use behaviors.

Lower segregation and higher integration, particularly during development, is suggestive of a more flexible and less specialized brain network^17^. This more “randomized”, less mature network topology in SCNs is often observed during typical neurodevelopment between the ages of 4 and 11, preceding the network specialization processes that occur later in adolescence^17,48^. While the long-term implications of the network differences observed in the current study remain unclear, it is worth noting that both protracted^49^ and accelerated^12,50^ markers of cortical development have been identified in early alcohol initiation. The network topology identified in our initiator group may therefore reflect a less mature network structure at this age (protracted) or an earlier shift towards the typical network changes that occur between ages 4 to 11 (accelerated). While the functional implications of these findings require further exploration, these results suggest atypical network properties in pre-adolescence precede early alcohol initiation, underscoring the importance of investigating the long-term whole-brain neuroanatomical mechanisms associated with early alcohol use.

Unlike previous studies using ABCD data^10,15^, the current study found no statistically significant regional predictors of early initiation of alcohol use. Importantly, the current study differed from these previous studies in terms of the definition of “initiation” of alcohol use. These previous studies defined alcohol initiation as either “sipping” *or* consuming a “full drink”. However, research has shown that early initiation of sipping may be more reflective of parenting practices and other external environmental influences, rather than a propensity to engage in problem behaviors that may be more reliably predicted by underlying biology^50^. As a result, our more stringent definition of initiation focused on consuming a full drink only. The lack of relationships between regional measures of brain structure and consuming a full drink of alcohol highlights the importance of modelling the brain’s network-level complexity when predicting more stringent conceptualizations of alcohol use initiation.

While these findings indicate novel neuroanatomical mechanisms associated with early alcohol use, these were only present in cortical thickness networks and not cortical volume networks. While cortical thickness is considered to capture meaningful biological variation associated with cognitive, behavioral, and neurodevelopmental changes^51^, these estimates are prone to relative measurement errors, and there have been concerns that these may yield unreliable estimates for SCN analyses^52^. To address this issue, cortical volume SCNs were also calculated to test for robustness. Despite a lack of statistical significance for the cortical volume network properties, the same direction of relationships were observed in the cortical volume SCNs, suggesting our results may reflect loss of sensitivity when using cortical volume^46^. These findings are also consistent with previous studies of regional neuroanatomical predictors of early alcohol use initiation, where regional cortical thickness was more likely to be related to early alcohol use initiation when compared with regional cortical volume^10,15^.

This study has several strengths, including the use of a large, community-based sample that broadly reflects the diversity of the US population, an alcohol-naïve sample to eliminate reverse causation, and a matched sample to reduce confounding bias. However, the stringent definition of alcohol initiation limited the sample size of initiators, potentially affecting the reliability of relationships between brain network organization and early alcohol use (reflected in effect sizes and variability in the network resilience measure). Finally, the small number of alcohol initiators reduced our capacity to examine other, more nuanced contributing factors linked to early alcohol initiation, such as peer influence^53^, family history of alcohol use^54^, and childhood trauma^55^, which is central to understanding the broader picture of early alcohol initiation.

In conclusion, we provide novel evidence to suggest that differences in structural brain network organization in alcohol-naïve adolescents precede early alcohol initiation prior to age 15, independent of regional differences in brain structure. Further research in a larger, longitudinal sample tracking potential psychosocial, behavioral, and environmental factors associated with these relationships would be beneficial.

### Data sharing statement

The data used in this study are from the Adolescent Brain Cognitive Development (ABCD) Study, an open-source dataset available to researchers. The ABCD Study data can be accessed via the National Institute of Mental Health Data Archive (NDA) at https://nda.nih.gov/abcd. Data sharing is subject to the terms and conditions outlined by the NDA, including registration and approval for access to the study’s datasets.

## Acknowledgements

Data used in the preparation of this article were obtained from the Adolescent Brain Cognitive Development^SM^ (ABCD) Study (https://abcdstudy.org), held in the NIMH Data Archive (NDA). This is a multisite, longitudinal study designed to recruit more than 10,000 children age 9-10 and follow them over 10 years into early adulthood. The ABCD Study® is supported by the National Institutes of Health and additional federal partners under award numbers U01DA041048, U01DA050989, U01DA051016, U01DA041022, U01DA051018, U01DA051037, U01DA050987, U01DA041174, U01DA041106, U01DA041117, U01DA041028, U01DA041134, U01DA050988, U01DA051039, U01DA041156, U01DA041025, U01DA041120, U01DA051038, U01DA041148, U01DA041093, U01DA041089, U24DA041123, U24DA041147. A full list of supporters is available at https://abcdstudy.org/federal-partners.html. A listing of participating sites and a complete listing of the study investigators can be found at https://abcdstudy.org/consortium_members/. ABCD consortium investigators designed and implemented the study and/or provided data but did not necessarily participate in the analysis or writing of this report. This manuscript reflects the views of the authors and may not reflect the opinions or views of the NIH or ABCD consortium investigators.

