## Supplementary tables and figures for "Structural Covariance Network Properties Predictive of Early Adolescent Alcohol Initiation"

**Supplementary Material**

**Network measures**

| **Network feature** | **Description** | **Network measure** | **Description** |
| --- | --- | --- | --- |
| Segregation | Measures of network segregation primarily quantify the presence of densely interconnected groups (known as clusters or modules) within the network, which enables specialized processing to occur. Higher network segregation is therefore considered to reflect greater functional specialization within the network. | Modularity | A measure of the presence of strongly interconnected subgroups |
|  |  | Clustering coefficient | A measure of clustering around individual nodes |
| Integration | Network integration describes the ability to rapidly combine information from distributed brain regions. Accordingly, higher network integration is typically associated with greater processing efficiency within the network. | Characteristic path length | The average shortest path length in the network |
|  |  | Global efficiency | The average of the inverse of the shortest path lengths in the network |
| Resilience | Network resilience quantifies features of the network that reflect vulnerability of the network to insult. Higher network resilience is generally associated with more efficient information processing and a lower vulnerability to network damage. | Degree assortativity | Correlation between the degrees of connected nodes in the network |

***eTable 1****.* Description of network measures used in the study, as described in Rubinov and Sporns (2010).

**Sample matching**

We used propensity score matching through the *MatchIt* package to balance covariates between groups. We first attempted 1:1 nearest neighbour propensity score matching without replacement with a propensity score estimated using logistic regression of group status on the covariates. This matching specification yielded poor balance, so we instead tried optimal matching based on the propensity score, which yielded adequate balance, as indicated in *eFigure 1*. After matching, all standardized mean differences for the covariates were below 0.1, indicating adequate balance for our study. Sample characteristics for the matched samples are listed in *eTable 2.*

***eFigure 1.*** Love plot showing covariate balance obtained from 1:1 optimal matching based on propensity scores using the *MatchIt* package.

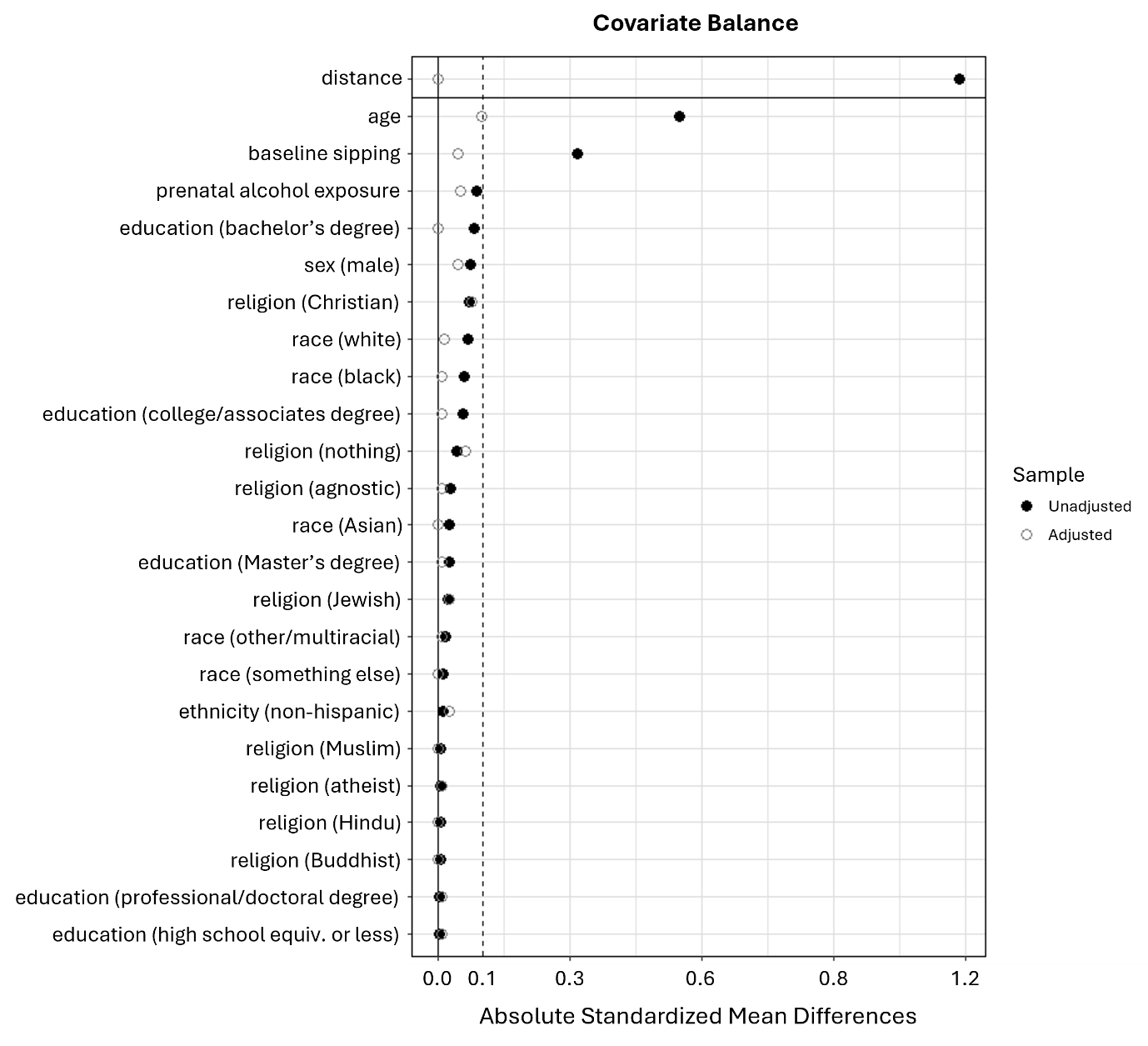

***eTable 2.*** Sample characteristics of the matched cohort.

| **Characteristic** | **Initiators (*N* = 160)** | **Non-Initiators (*N* = 160)** | ***p*-value** |
| --- | --- | --- | --- |
| **Baseline** |  |  |  |
| Age at baseline (years)^1,3^ | 10.4 (10-10.8) | 10.3 (9.8-10.8) | 0.3 |
| Sex^2,4^ |  |  | 0.4 |
| Female | 87 (54.4%) | 94 (58.8%) |  |
| Race^2,5^ |  |  | >0.9 |
| Asian | 0 | 0 |  |
| Black/African American, Other/Multiracial^†^ | 35 (21.9%) | 37 (23.1%) |  |
| White | 125 (78.1%) | 123 (76.9%) |  |
| Ethnicity^2,4^ |  |  | 0.6 |
| Hispanic/Latino | 34 (21.3%) | 38 (23.8%) |  |
| Religion^2,5^ |  |  | 0.7 |
| Christian | 95 (59.4%) | 107 (66.9%) |  |
| Jewish, Muslim, Buddhist, Hindu, Something else^†^ | 9 (5.6%) | 5 (3.1%) |  |
| Atheist, Agnostic^†^ | 13 (8.1%) | 15 (9.4%) |  |
| Nothing | 43 (26.9%) | 33 (20.6%) |  |
| Highest Parental Education^2,4^ |  |  | 0.2 |
| High School | 21 (13.1%) | 20 (12.5%) |  |
| College/Associate’s Degree | 54 (33.8%) | 55 (34.4%) |  |
| Bachelor’s Degree | 37 (23.1%) | 37 (23.1%) |  |
| Master’s Degree | 37 (23.1%) | 38 (23.8%) |  |
| Professional/Doctoral Degree | 11 (6.9%) | 10 (6.3%) |  |
| Prenatal alcohol exposure^2,4^ | 58 (36.3%) | 66 (41.3%) | 0.4 |
| Alcohol sipping at baseline^2,4^ | 89 (55.6%) | 96 (60%) | 0.4 |

^1^ Median (IQR); ^2^ n (%); ^3^ Wilcoxon rank sum test; ^4^ Pearson’s Chi-squared test; ^5^ Fisher’s exact test. ^†^ Categories have been collapsed for reporting purposes due to low sample sizes and possible identifiability. However, analyses were conducted with all levels of each category preserved.

**Mean cortical thickness adjustment**

While whole brain volume demonstrates a strong linear relationship with regional cortical and subcortical volumes (Jäncke et al., 2015), the question of whether to adjust for mean cortical thickness when comparing regional cortical thickness measurements between groups is currently under debate. This is due in large part to the considerable variability in regional cortical thickness that is typically present across the cortex, thus adjusting for whole-brain measures has the potential to obscure meaningful inter-individual differences (Vijayakumar et al., 2018; Backhausen et al., 2022). Therefore, linear models examining the association between alcohol use initiation (initiators vs. non-initiators) and cortical thickness measurements per hemisphere controlling for age, sex and mean global cortical thickness were performed, with output coefficients examined to determine how much variance within the models could be explained by mean cortical thickness. As demonstrated in *eTable 3*, results from the linear models revealed that mean cortical thickness accounted for a meaningful proportion of variance in regional cortical thickness across 21/68 ROIs (*R*^2^ > 0.4) and was significantly associated with the outcome across all 68 ROIs (all p_FDR_ values < .001). Based on these findings, mean global cortical thickness was incorporated as a first and second level covariate in all statistical models comparing cortical thickness measurements, including the subsequent SCN analyses.

|  | **Left hemisphere** | | | | | | **Right hemisphere** | | | | | |
| --- | --- | --- | --- | --- | --- | --- | --- | --- | --- | --- | --- | --- |
| **Region** | **Estimate** | **SE** | ***t*** | ***P*** | ***P*_FDR_** | ***R^2^*** | **Estimate** | **SE** | ***t*** | ***P*** | ***P*_FDR_** | ***R^2^*** |
| BSTS | 0.992 | 0.028 | 35.324 | <.0001*** | <.0001*** | 0.251 | 1.037 | 0.029 | 36.174 | <.0001*** | <.0001*** | 0.256 |
| cACC | 0.490 | 0.038 | 12.910 | <.0001*** | <.0001*** | 0.053 | 0.466 | 0.036 | 12.985 | <.0001*** | <.0001*** | 0.046 |
| cMFG | 1.054 | 0.020 | 52.198 | <.0001*** | <.0001*** | 0.420 | 1.029 | 0.021 | 50.081 | <.0001*** | <.0001*** | 0.401 |
| CUN | 0.887 | 0.024 | 36.276 | <.0001*** | <.0001*** | 0.268 | 0.871 | 0.023 | 37.226 | <.0001*** | <.0001*** | 0.276 |
| ENT | 0.995 | 0.053 | 18.926 | <.0001*** | <.0001*** | 0.085 | 1.013 | 0.058 | 17.372 | <.0001*** | <.0001*** | 0.079 |
| FUS | 0.935 | 0.017 | 54.991 | <.0001*** | <.0001*** | 0.445 | 0.926 | 0.017 | 53.069 | <.0001*** | <.0001*** | 0.427 |
| IPL | 1.013 | 0.016 | 64.703 | <.0001*** | <.0001*** | 0.529 | 1.014 | 0.015 | 66.749 | <.0001*** | <.0001*** | 0.542 |
| ITG | 1.035 | 0.021 | 49.390 | <.0001*** | <.0001*** | 0.389 | 1.038 | 0.021 | 50.304 | <.0001*** | <.0001*** | 0.397 |
| iCC | 0.499 | 0.028 | 17.768 | <.0001*** | <.0001*** | 0.080 | 0.422 | 0.029 | 14.589 | <.0001*** | <.0001*** | 0.058 |
| LOG | 0.996 | 0.017 | 59.834 | <.0001*** | <.0001*** | 0.484 | 1.046 | 0.017 | 61.395 | <.0001*** | <.0001*** | 0.497 |
| LOF | 0.833 | 0.020 | 42.643 | <.0001*** | <.0001*** | 0.329 | 0.808 | 0.021 | 38.588 | <.0001*** | <.0001*** | 0.285 |
| LING | 0.868 | 0.021 | 41.270 | <.0001*** | <.0001*** | 0.320 | 0.841 | 0.020 | 41.801 | <.0001*** | <.0001*** | 0.326 |
| MOF | 0.622 | 0.024 | 25.864 | <.0001*** | <.0001*** | 0.160 | 0.675 | 0.022 | 30.273 | <.0001*** | <.0001*** | 0.207 |
| MTG | 1.165 | 0.024 | 49.437 | <.0001*** | <.0001*** | 0.391 | 1.153 | 0.021 | 54.095 | <.0001*** | <.0001*** | 0.434 |
| PARH | 0.857 | 0.049 | 17.630 | <.0001*** | <.0001*** | 0.096 | 0.841 | 0.040 | 20.964 | <.0001*** | <.0001*** | 0.132 |
| paraC | 1.095 | 0.021 | 51.257 | <.0001*** | <.0001*** | 0.414 | 1.076 | 0.020 | 52.757 | <.0001*** | <.0001*** | 0.426 |
| pOPER | 0.897 | 0.020 | 44.820 | <.0001*** | <.0001*** | 0.344 | 0.851 | 0.022 | 38.316 | <.0001*** | <.0001*** | 0.280 |
| pORB | 1.082 | 0.031 | 35.254 | <.0001*** | <.0001*** | 0.258 | 1.022 | 0.030 | 34.411 | <.0001*** | <.0001*** | 0.253 |
| pTRI | 0.934 | 0.023 | 41.438 | <.0001*** | <.0001*** | 0.317 | 0.902 | 0.021 | 42.859 | <.0001*** | <.0001*** | 0.327 |
| periCAL | 0.831 | 0.026 | 32.198 | <.0001*** | <.0001*** | 0.216 | 0.822 | 0.027 | 30.465 | <.0001*** | <.0001*** | 0.196 |
| postC | 1.036 | 0.022 | 47.557 | <.0001*** | <.0001*** | 0.373 | 1.004 | 0.024 | 41.397 | <.0001*** | <.0001*** | 0.316 |
| PCC | 0.528 | 0.025 | 21.533 | <.0001*** | <.0001*** | 0.113 | 0.502 | 0.023 | 22.136 | <.0001*** | <.0001*** | 0.117 |
| preC | 1.107 | 0.019 | 57.924 | <.0001*** | <.0001*** | 0.470 | 1.069 | 0.023 | 46.648 | <.0001*** | <.0001*** | 0.365 |
| PCUN | 0.978 | 0.014 | 68.575 | <.0001*** | <.0001*** | 0.558 | 0.985 | 0.014 | 68.276 | <.0001*** | <.0001*** | 0.557 |
| rACC | 0.643 | 0.034 | 18.790 | <.0001*** | <.0001*** | 0.101 | 0.595 | 0.036 | 16.567 | <.0001*** | <.0001*** | 0.080 |
| rMFG | 0.978 | 0.016 | 61.956 | <.0001*** | <.0001*** | 0.503 | 0.949 | 0.016 | 58.612 | <.0001*** | <.0001*** | 0.477 |
| SFG | 1.133 | 0.016 | 70.755 | <.0001*** | <.0001*** | 0.570 | 1.133 | 0.016 | 70.452 | <.0001*** | <.0001*** | 0.570 |
| SPL | 1.023 | 0.014 | 70.564 | <.0001*** | <.0001*** | 0.568 | 1.024 | 0.015 | 70.137 | <.0001*** | <.0001*** | 0.565 |
| STG | 1.197 | 0.022 | 55.034 | <.0001*** | <.0001*** | 0.445 | 1.195 | 0.020 | 58.995 | <.0001*** | <.0001*** | 0.476 |
| SMAR | 1.120 | 0.019 | 59.744 | <.0001*** | <.0001*** | 0.486 | 1.076 | 0.018 | 58.265 | <.0001*** | <.0001*** | 0.471 |
| FP | 1.104 | 0.048 | 23.189 | <.0001*** | <.0001*** | 0.128 | 1.118 | 0.047 | 23.982 | <.0001*** | <.0001*** | 0.135 |
| TP | 1.371 | 0.050 | 27.499 | <.0001*** | <.0001*** | 0.164 | 1.323 | 0.051 | 26.144 | <.0001*** | <.0001*** | 0.154 |
| TT | 1.172 | 0.033 | 35.435 | <.0001*** | <.0001*** | 0.264 | 1.122 | 0.034 | 33.008 | <.0001*** | <.0001*** | 0.228 |
| INS | 0.707 | 0.026 | 26.884 | <.0001*** | <.0001*** | 0.167 | 0.711 | 0.027 | 26.092 | <.0001*** | <.0001*** | 0.160 |

***eTable 3.*** Results from the linear model comparing cortical thickness measurements between groups while controlling for age, sex, and mean cortical thickness. Results show the output coefficients from the mean cortical thickness variable within the linear model.

|  | | **Thickness** | | | | | | **Volume** | | | | | |
| --- | --- | --- | --- | --- | --- | --- | --- | --- | --- | --- | --- | --- | --- |
| **Hem** | **Region** | **Estimate** | **SE** | ***t*** | ***P*** | ***P*_FDR_** | ***R^2^*** | **Estimate** | **SE** | ***t*** | ***P*** | ***P*_FDR_** | ***R^2^*** |
| Left | BSTS | 0.005 | 0.009 | 0.544 | 0.586 | 0.919 | 0.251 | -23.552 | 37.442 | -0.629 | 0.529 | 0.886 | 0.210 |
|  | cACC | -0.002 | 0.012 | -0.170 | 0.865 | 0.962 | 0.053 | -55.348 | 36.872 | -1.501 | 0.133 | 0.831 | 0.138 |
|  | cMFG | 0.000 | 0.007 | -0.038 | 0.970 | 0.970 | 0.420 | -5.615 | 88.317 | -0.064 | 0.949 | 0.982 | 0.276 |
|  | CUN | 0.002 | 0.008 | 0.211 | 0.833 | 0.962 | 0.268 | 59.415 | 42.710 | 1.391 | 0.164 | 0.831 | 0.164 |
|  | ENT | -0.004 | 0.017 | -0.242 | 0.809 | 0.962 | 0.085 | 9.172 | 26.875 | 0.341 | 0.733 | 0.923 | 0.141 |
|  | FUS | -0.007 | 0.005 | -1.338 | 0.181 | 0.810 | 0.445 | -109.045 | 81.042 | -1.346 | 0.179 | 0.831 | 0.433 |
|  | IPL | 0.006 | 0.005 | 1.158 | 0.247 | 0.810 | 0.529 | 56.773 | 136.428 | 0.416 | 0.677 | 0.886 | 0.304 |
|  | ITG | 0.002 | 0.007 | 0.246 | 0.805 | 0.962 | 0.389 | -50.024 | 108.298 | -0.462 | 0.644 | 0.886 | 0.401 |
|  | iCC | 0.001 | 0.009 | 0.094 | 0.925 | 0.970 | 0.080 | -1.022 | 30.071 | -0.034 | 0.973 | 0.982 | 0.343 |
|  | LOG | -0.003 | 0.005 | -0.532 | 0.595 | 0.919 | 0.484 | -14.649 | 110.965 | -0.132 | 0.895 | 0.982 | 0.373 |
|  | LOF | 0.002 | 0.006 | 0.245 | 0.807 | 0.962 | 0.329 | -12.271 | 52.609 | -0.233 | 0.816 | 0.982 | 0.488 |
|  | LING | -0.006 | 0.007 | -0.887 | 0.375 | 0.810 | 0.320 | 35.785 | 74.291 | 0.482 | 0.630 | 0.886 | 0.221 |
|  | MOF | 0.017 | 0.008 | 2.225 | **0.026*** | 0.603 | 0.160 | 25.000 | 41.121 | 0.608 | 0.543 | 0.886 | 0.402 |
|  | MTG | 0.003 | 0.008 | 0.441 | 0.659 | 0.933 | 0.391 | 151.037 | 100.090 | 1.509 | 0.131 | 0.831 | 0.428 |
|  | PARH | -0.001 | 0.016 | -0.070 | 0.944 | 0.970 | 0.096 | -21.348 | 22.663 | -0.942 | 0.346 | 0.831 | 0.150 |
|  | paraC | 0.008 | 0.007 | 1.169 | 0.243 | 0.810 | 0.414 | -0.803 | 35.834 | -0.022 | 0.982 | 0.982 | 0.255 |
|  | pOPER | 0.005 | 0.006 | 0.736 | 0.461 | 0.848 | 0.344 | -81.872 | 65.184 | -1.256 | 0.209 | 0.831 | 0.188 |
|  | pORB | -0.012 | 0.010 | -1.170 | 0.242 | 0.810 | 0.258 | -26.409 | 24.947 | -1.059 | 0.290 | 0.831 | 0.279 |
|  | pTRI | 0.007 | 0.007 | 0.952 | 0.341 | 0.810 | 0.317 | -22.612 | 52.452 | -0.431 | 0.666 | 0.886 | 0.183 |
|  | periCAL | -0.004 | 0.008 | -0.480 | 0.631 | 0.933 | 0.216 | 37.390 | 31.108 | 1.202 | 0.229 | 0.831 | 0.114 |
|  | postC | -0.005 | 0.007 | -0.728 | 0.467 | 0.848 | 0.373 | 8.563 | 92.720 | 0.092 | 0.926 | 0.982 | 0.391 |
|  | PCC | 0.010 | 0.008 | 1.318 | 0.188 | 0.810 | 0.113 | -22.880 | 36.218 | -0.632 | 0.528 | 0.886 | 0.307 |
|  | preC | 0.010 | 0.006 | 1.570 | 0.117 | 0.810 | 0.470 | -58.937 | 100.369 | -0.587 | 0.557 | 0.886 | 0.422 |
|  | PCUN | -0.006 | 0.005 | -1.344 | 0.179 | 0.810 | 0.558 | -68.848 | 86.371 | -0.797 | 0.425 | 0.851 | 0.439 |
|  | rACC | 0.010 | 0.011 | 0.876 | 0.381 | 0.810 | 0.101 | -38.638 | 35.575 | -1.086 | 0.278 | 0.831 | 0.313 |
|  | rMFG | 0.005 | 0.005 | 0.916 | 0.360 | 0.810 | 0.503 | -76.599 | 142.352 | -0.538 | 0.591 | 0.886 | 0.404 |
|  | SFG | 0.003 | 0.005 | 0.637 | 0.524 | 0.891 | 0.570 | 149.619 | 174.389 | 0.858 | 0.391 | 0.831 | 0.477 |
|  | SPL | -0.003 | 0.005 | -0.716 | 0.474 | 0.848 | 0.568 | 6.802 | 138.278 | 0.049 | 0.961 | 0.982 | 0.315 |
|  | STG | -0.007 | 0.007 | -1.057 | 0.291 | 0.810 | 0.445 | 211.175 | 104.420 | 2.022 | **0.043*** | 0.831 | 0.430 |
|  | SMAR | -0.013 | 0.006 | -2.104 | **0.035*** | 0.603 | 0.486 | 138.538 | 146.459 | 0.946 | 0.344 | 0.831 | 0.339 |
|  | FP | 0.002 | 0.015 | 0.155 | 0.877 | 0.962 | 0.128 | 10.886 | 12.181 | 0.894 | 0.372 | 0.831 | 0.193 |
|  | TP | -0.020 | 0.016 | -1.241 | 0.215 | 0.810 | 0.164 | -25.868 | 24.779 | -1.044 | 0.297 | 0.831 | 0.169 |
|  | TT | -0.011 | 0.011 | -1.032 | 0.302 | 0.810 | 0.264 | 20.817 | 16.411 | 1.268 | 0.205 | 0.831 | 0.201 |
|  | INS | 0.003 | 0.008 | 0.372 | 0.710 | 0.962 | 0.167 | 50.450 | 45.228 | 1.115 | 0.265 | 0.831 | 0.451 |
| Right | BSTS | -0.004 | 0.009 | -0.387 | 0.699 | 0.973 | 0.256 | -4.405 | 28.937 | -0.152 | 0.879 | 0.966 | 0.263 |
|  | cACC | -0.001 | 0.012 | -0.115 | 0.909 | 0.973 | 0.046 | -3.640 | 39.129 | -0.093 | 0.926 | 0.966 | 0.130 |
|  | cMFG | -0.001 | 0.007 | -0.212 | 0.832 | 0.973 | 0.401 | 11.523 | 89.314 | 0.129 | 0.897 | 0.966 | 0.241 |
|  | CUN | -0.003 | 0.008 | -0.390 | 0.696 | 0.973 | 0.276 | -20.856 | 44.301 | -0.471 | 0.638 | 0.966 | 0.198 |
|  | ENT | -0.005 | 0.019 | -0.287 | 0.774 | 0.973 | 0.079 | -6.475 | 26.694 | -0.243 | 0.808 | 0.966 | 0.115 |
|  | FUS | 0.003 | 0.006 | 0.457 | 0.648 | 0.973 | 0.427 | -14.480 | 75.437 | -0.192 | 0.848 | 0.966 | 0.477 |
|  | IPL | -0.007 | 0.005 | -1.348 | 0.178 | 0.958 | 0.542 | -74.768 | 157.704 | -0.474 | 0.635 | 0.966 | 0.386 |
|  | ITG | -0.009 | 0.007 | -1.295 | 0.195 | 0.958 | 0.397 | -130.144 | 100.771 | -1.291 | 0.197 | 0.966 | 0.416 |
|  | iCC | 0.010 | 0.009 | 1.083 | 0.279 | 0.958 | 0.058 | 14.311 | 29.043 | 0.493 | 0.622 | 0.966 | 0.281 |
|  | LOG | 2.46E-05 | 0.006 | 0.004 | 0.996 | 0.996 | 0.497 | 19.470 | 118.945 | 0.164 | 0.870 | 0.966 | 0.382 |
|  | LOF | -0.006 | 0.007 | -0.902 | 0.367 | 0.958 | 0.285 | -38.077 | 53.105 | -0.717 | 0.473 | 0.966 | 0.485 |
|  | LING | 0.001 | 0.006 | 0.139 | 0.889 | 0.973 | 0.326 | 5.960 | 75.692 | 0.079 | 0.937 | 0.966 | 0.237 |
|  | MOF | 0.007 | 0.007 | 0.996 | 0.319 | 0.958 | 0.207 | -14.658 | 41.250 | -0.355 | 0.722 | 0.966 | 0.443 |
|  | MTG | -0.007 | 0.007 | -1.067 | 0.286 | 0.958 | 0.434 | 29.869 | 100.144 | 0.298 | 0.766 | 0.966 | 0.475 |
|  | PARH | -0.002 | 0.013 | -0.173 | 0.863 | 0.973 | 0.132 | -15.352 | 18.891 | -0.813 | 0.416 | 0.966 | 0.183 |
|  | paraC | -0.001 | 0.007 | -0.192 | 0.847 | 0.973 | 0.426 | 34.374 | 41.805 | 0.822 | 0.411 | 0.966 | 0.254 |
|  | pOPER | 0.003 | 0.007 | 0.428 | 0.669 | 0.973 | 0.280 | -34.395 | 51.481 | -0.668 | 0.504 | 0.966 | 0.202 |
|  | pORB | -0.006 | 0.010 | -0.647 | 0.518 | 0.973 | 0.253 | -0.969 | 29.992 | -0.032 | 0.974 | 0.974 | 0.258 |
|  | pTRI | -0.001 | 0.007 | -0.086 | 0.931 | 0.973 | 0.327 | -70.569 | 60.546 | -1.166 | 0.244 | 0.966 | 0.188 |
|  | periCAL | 0.002 | 0.009 | 0.282 | 0.778 | 0.973 | 0.196 | 17.379 | 32.599 | 0.533 | 0.594 | 0.966 | 0.136 |
|  | postC | 0.007 | 0.008 | 0.853 | 0.394 | 0.958 | 0.316 | 67.487 | 90.264 | 0.748 | 0.455 | 0.966 | 0.373 |
|  | PCC | -0.001 | 0.007 | -0.202 | 0.840 | 0.973 | 0.117 | 12.540 | 36.539 | 0.343 | 0.731 | 0.966 | 0.311 |
|  | preC | -0.010 | 0.007 | -1.362 | 0.173 | 0.958 | 0.365 | -75.351 | 104.086 | -0.724 | 0.469 | 0.966 | 0.385 |
|  | PCUN | 0.004 | 0.005 | 0.852 | 0.395 | 0.958 | 0.557 | -65.445 | 87.971 | -0.744 | 0.457 | 0.966 | 0.462 |
|  | rACC | -0.001 | 0.012 | -0.070 | 0.944 | 0.973 | 0.080 | 5.413 | 30.644 | 0.177 | 0.860 | 0.966 | 0.205 |
|  | rMFG | 0.007 | 0.005 | 1.299 | 0.194 | 0.958 | 0.477 | -21.783 | 158.295 | -0.138 | 0.891 | 0.966 | 0.365 |
|  | SFG | -0.001 | 0.005 | -0.171 | 0.864 | 0.973 | 0.570 | 321.049 | 175.473 | 1.830 | 0.067 | 0.764 | 0.462 |
|  | SPL | -0.006 | 0.005 | -1.350 | 0.177 | 0.958 | 0.565 | -148.408 | 137.724 | -1.078 | 0.281 | 0.966 | 0.334 |
|  | STG | 0.001 | 0.007 | 0.079 | 0.937 | 0.973 | 0.476 | 205.867 | 89.827 | 2.292 | **0.022*** | 0.373 | 0.430 |
|  | SMAR | 0.002 | 0.006 | 0.383 | 0.701 | 0.973 | 0.471 | -9.507 | 120.869 | -0.079 | 0.937 | 0.966 | 0.322 |
|  | FP | 0.022 | 0.015 | 1.474 | 0.141 | 0.958 | 0.135 | 38.392 | 14.477 | 2.652 | **0.008*** | 0.273 | 0.180 |
|  | TP | 0.014 | 0.016 | 0.874 | 0.382 | 0.958 | 0.154 | -3.046 | 25.038 | -0.122 | 0.903 | 0.966 | 0.151 |
|  | TT | -0.004 | 0.011 | -0.370 | 0.711 | 0.973 | 0.228 | 12.771 | 11.681 | 1.093 | 0.274 | 0.966 | 0.242 |
|  | INS | 0.009 | 0.009 | 1.058 | 0.290 | 0.958 | 0.160 | 70.721 | 43.777 | 1.615 | 0.106 | 0.903 | 0.452 |

***eTable 4.*** Results from linear models comparing cortical thickness and cortical volume measurements between groups controlling for age, sex, and whole-brain measurements (mean cortical thickness and eTIV, respectively).

**BSTS** = bank of the superior temporal sulcus; **cACC** = caudal anterior cingulate; **cMFG** = caudal middle frontal gyrus; **CUN** = cuneus; **ENT** = entorhinal; **FUS** = fusiform; **IPL** = inferior parietal lobule; **ITG** = inferior temporal gyrus; **iCC** = isthmus cingulate cortex; **LOG** = lateral occipital gyrus; **LOF** = lateral orbitofrontal; **LING** = lingual; **MOF** = medial orbitofrontal; **MTG** = middle temporal gyrus; **PARH** = parahippocampal; **paraC** = paracentral; **pOPER** = pars opercularis; **pORB** = pars orbitalis; **pTRI** = pars triangularis; **periCAL** = pericalcarine; **postC** = postcentral; **PCC** = posterior cingulate cortex; **preC** = precentral; **PCUN** = precuneus; **rACC** = rostral anterior cingulate cortex; **rMFG** = rostral middle frontal gyrus; **SFG** = superior frontal gyrus; **SPL** = superior parietal lobule; **STG** = superior temporal gyrus; **SMAR** = supramarginal gyrus; **FP** = frontal pole; **TP** = temporal pole; **TT** = transverse temporal; **INS** = insula.

**Sensitivity analyses**

*SCN comparisons with second-level covariate adjustment*

When investigating group differences in cortical thickness structural covariance network measures adjusting for second-level covariates (age, sex, race, ethnicity, religion, parental education, prenatal alcohol exposure, and baseline alcohol sipping), the initiator group exhibited lower characteristic path length (AUC [95% CI] = -0.082 [-0.107, -0.049], p = 0.010) and degree assortativity (AUC [95% CI] = -0.024 [-0.047, -0.002], p = 0.040), in addition to higher global efficiency (AUC [95% CI] = 0.008 [0.006, 0.010], p = 0.0495), compared to non-initiators. When examining the cortical volume structural covariance networks, there were no statistically significant AUC differences in network measures between initiators and non-initiators were identified. Overall, these findings replicate the relationships observed when comparing against the matched samples.

*Other substance use initiation in control group*

As individuals who initiated use of other substances during follow-up may present different network properties at baseline compared to total abstainers, a sensitivity analysis was performed for the cortical thickness SCN analysis excluding participants in the non-initiator group who initiated other substance use, defined as cigarette, e-cigarette, or marijuana use (n=142). When investigating group differences in cortical thickness structural covariance graph-level network properties controlling for first-level covariates, the initiator group exhibited lower characteristic path length (AUC [95% CI] = -0.069 [-0.108, -0.036], p = 0.0495) and degree assortativity (AUC [95% CI] = -0.024 [-0.050, 0.004], p = 0.020), in addition to higher global efficiency (AUC [95% CI] = 0.008 [0.005, 0.010], p = 0.010), compared to non-initiators, replicating the findings observed in the full sample. These findings were replicated in the matched sample (characteristic path length: AUC [95% CI] = -0.001 [-0.032, 0.040], p = 0.0495; degree assortativity: AUC [95% CI] = -0.001 [-0.028, 0.027], p = 0.040; global efficiency: AUC [95% CI] = 0.001 [-0.003, 0.003], p = 0.040) and when adjusting for second-level covariates (characteristic path length: AUC [95% CI] = -0.069 [-0.101, -0.037], p = 0.030; degree assortativity: AUC [95% CI] = -0.026 [-0.047, -0.004], p = 0.020; global efficiency: AUC [95% CI] = 0.008 [0.005, 0.010], p = 0.010).

**Post hoc tests**

*Bootstrapping*

In the full sample cortical thickness SCN comparisons, the 95% CIs for degree assortativity crossed zero, indicating uncertainty in the magnitude of the observed effects. This was also observed in the matched sample for characteristic path length, degree assortativity, and global efficiency. To explore whether this uncertainty was due to instability of these network measures across different densities, bootstrapping with 1,000 iterations was performed using *brainGraph* (Watson, 2016). For degree assortativity, in both the full and matched samples, the bootstrapping results indicated CIs were wide for both groups, especially for the initiator group at lower densities (*eFigs. 2* and *3*), suggesting low stability may explain the crossing of CIs observed in the degree assortativity AUC tests. In contrast, for characteristic path length and global efficiency in the matched sample, CIs for both groups were narrow, suggesting CI crossing in the AUC tests for these measures are likely unrelated to stability across densities and may be related to other factors, such as small sample size.

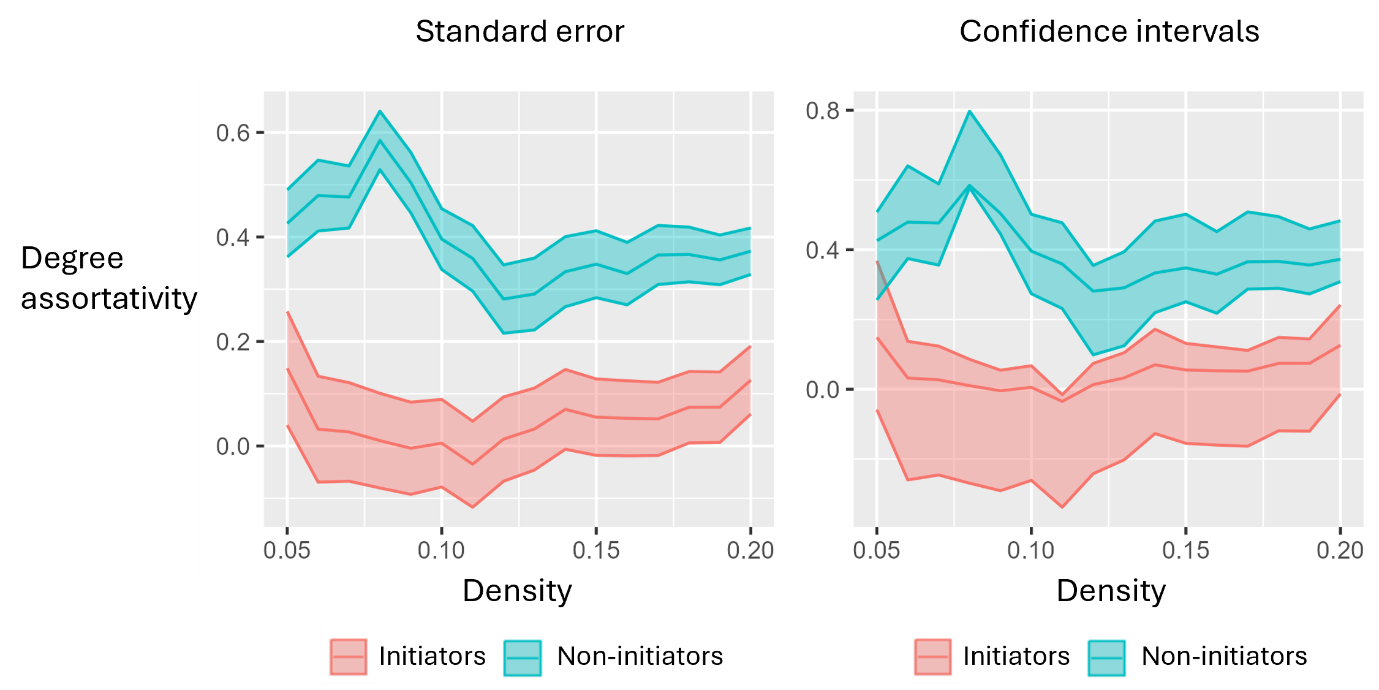

**eFigure 2.** Plots showing bootstrapping results for SCN degree assortativity between groups for the full and matched samples. Plots in the left column shows the ±1 standard error across densities for each measure, the right column displays the 95% confidence intervals.

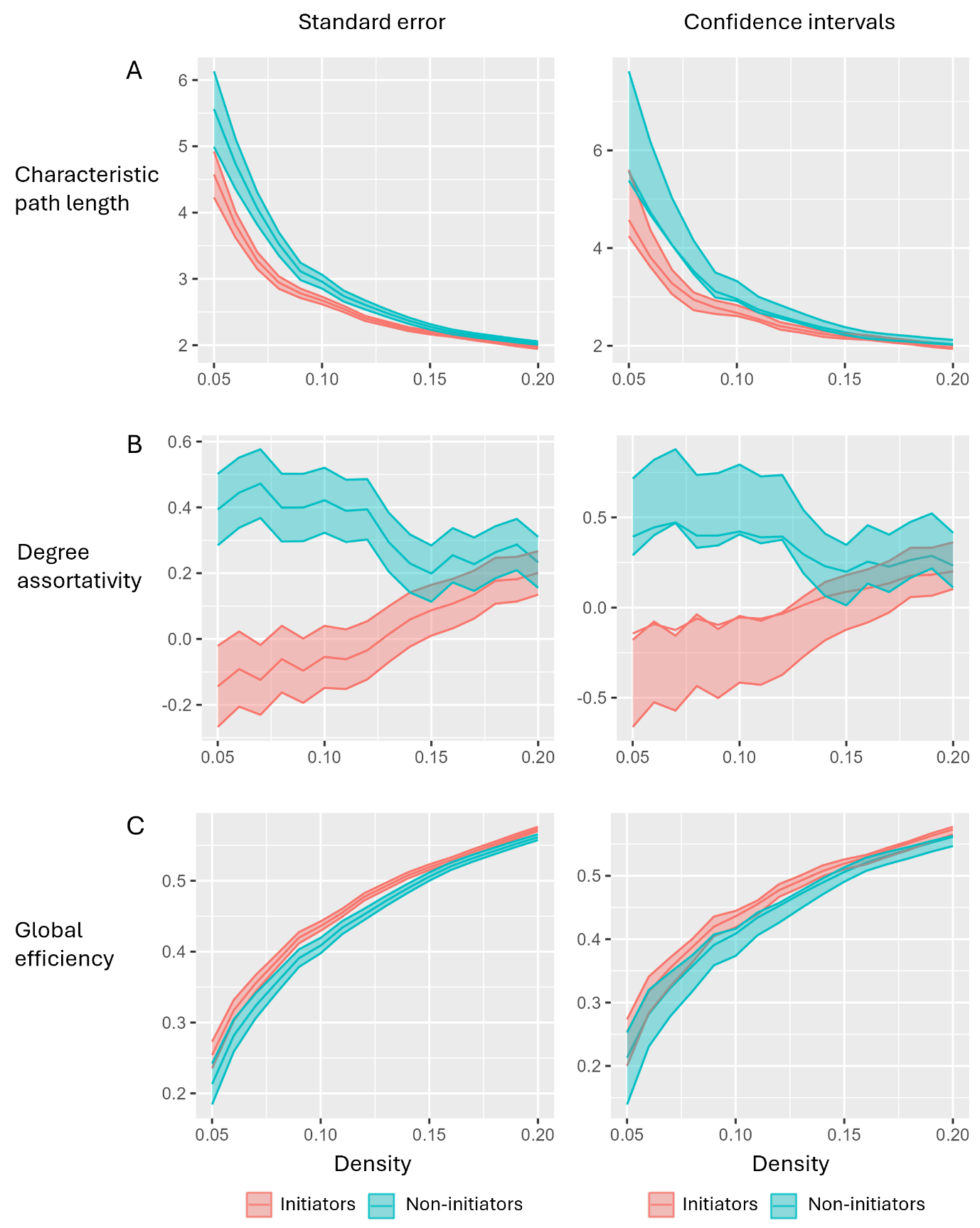

**eFigure 3.** Plots showing bootstrapping results for the matched samples cortical thickness SCN measures, including (A) characteristic path length; (B) degree assortativity, and; (C) global efficiency. Plots in the left column shows the ±1 standard error across densities for each measure, the right column displays the 95% confidence intervals.
